# Whole blood immunophenotyping uncovers immature neutrophil-to-VD2 T-cell ratio as an early prognostic marker for severe COVID-19

**DOI:** 10.1101/2020.06.11.147389

**Authors:** Guillaume Carissimo, Weili Xu, Immanuel Kwok, Mohammad Yazid Abdad, Yi-Hao Chan, Siew-Wai Fong, Kia Joo Puan, Cheryl Yi-Pin Lee, Nicholas Kim-Wah Yeo, Siti Naqiah Amrun, Rhonda Sin-Ling Chee, Wilson How, Stephrene Chan, Eugene Bingwen Fan, Anand Kumar Andiappan, Bernett Lee, Olaf Rötzschke, Barnaby Edward Young, Yee-Sin Leo, David C. Lye, Laurent Renia, Lai Guan Ng, Anis Larbi, Lisa F.P. Ng

**Affiliations:** Singapore Immunology Network, Agency for Science, Technology and Research, Immunos, Biopolis, 138648, Singapore; National Centre for Infectious Diseases, 16 Jalan Tan Tock Seng, 308442, Singapore; Department of Biological Sciences, National University of Singapore, Singapore 117543; Department of Haematology, Tan Tock Seng Hospital, 11 Jalan Tan Tock Seng, 308433, Singapore; Department of Infectious Diseases, Tan Tock Seng Hospital, 11 Jalan Tan Tock Seng, 308433, Singapore; Lee Kong Chian School of Medicine, Nanyang Technological University, 11 Mandalay Road, 308232, Singapore; Yong Loo Lin School of Medicine, National University of Singapore and National University Health System, 10 Medical Drive, 117597, Singapore; Department of Biochemistry, Yong Loo Lin School of Medicine, National University of Singapore, 8 Medical Drive, 117596, Singapore; Institute of Infection, Veterinary and Ecological Sciences, University of Liverpool, Liverpool, 8 West Derby Street, Liverpool L7 3EA, United Kingdom

**Keywords:** COVID-19, SARS-CoV-2, flow cytometry, immunophenotyping, monocytes, neutrophils, lymphocytes, gamma delta T-cells, cytokine, patients whole blood phenotyping

## Abstract

SARS-CoV-2 is the novel coronavirus responsible for the current COVID-19 pandemic. Severe complications are observed only in a small proportion of infected patients but the cellular mechanisms underlying this progression are still unknown. Comprehensive flow cytometry of whole blood samples from 54 COVID-19 patients revealed a dramatic increase in the number of immature neutrophils. This increase strongly correlated with disease severity and was associated with elevated IL-6 and IP-10 levels, two key players in the cytokine storm. The most pronounced decrease in cell counts was observed for CD8 T-cells and VD2 γδ T-cells, which both exhibited increased differentiation and activation. ROC analysis revealed that the count ratio of immature neutrophils to CD8 or VD2 T-cells predicts pneumonia onset (0.9071) as well as hypoxia onset (0.8908) with high sensitivity and specificity. It would thus be a useful prognostic marker for preventive patient management and improved healthcare resource management.

## Introduction

Severe Acute Respiratory Syndrome coronavirus 2 (SARS-CoV-2) first appeared in Wuhan, China in late 2019. It is a novel pathogen responsible for the coronavirus disease 2019 (COVID-19) pandemic ^1^. COVID-19 patients experience a wide spectrum of clinical manifestations that ranges from low-grade fever and mild respiratory symptoms, to more severe forms. This including acute respiratory distress syndrome (ARDS), which requires provision of supplemental oxygen, and in some cases intubation and mechanical ventilation ^2-5^. However, it remains unclear how SARS-CoV-2 infection affects the activation of immune cells and their contribution towards the severity of disease outcomes in patients.

Previous clinical studies reported associations with clinical blood counts, while others have specifically assessed T-cell subsets for activation and exhaustion markers ^6-9^. Since strong evidence points to a cytokine storm as the culprit for disease severity ^10,11^, various groups have investigated cytokine-secreting pathogenic T-cells and inflammatory monocytes that could have triggered this phenomenon ^6-9^. In addition, flow cytometry analysis in COVID-19 patients has also shown a polarisation towards the Th17 subtype and a highly activated and exhausted CD8^+^ T-cell compartment ^12,13^. All these stuies were carried out on peripheral blood mononuclear cells (PBMCs), thus excluding most granulocyte populations ^12,13^. However, to elucidate all the immune subsets that could potentially trigger severe COVID-19 pathology, it is imperative to perform comprehensive whole blood immunophenotyping of COVID-19 patients which includes granulocyte populations.

In this study, we employed high dimensional flow cytometry to analyse a wide spectrum of more than 50 subsets of the myeloid and lymphoid immune cell compartments. The study was carried out during the ongoing SARS-CoV-2 pandemic in Singapore with a cohort of 54 COVID-19 patients who presented with varied clinical manifestations ranging from mild to fatal outcomes. This comprehensive immunophenotyping allowed the identification of immature neutrophils, CD8 T-cells and gamma delta (VD) 2 T-cells as key immune cell populations that undergo substantial changes in the cell counts across the spectrum of clinical severity. Their numbers, in fact, represent an early and robust prognosis value as shown by ‘receiver operating characteristics’ (ROC) analysis.

## Results

### Circulating myeloid populations are reduced in COVID-19 patients

A total of 54 patients with laboratory-confirmed SARS-CoV-2 infection were recruited at the National Centre for Infectious Diseases (NCID), Singapore from end March to mid-May 2020 (Supplementary Table 1). Blood was collected from 54 patients upon enrollment at a median 7 days post-illness onset (pio), from 28 patients who had recovered from COVID-19 disease (median 30 days pio, Supplementary Table 1) and 19 healthy donors (Supplementary Table 2). Immunophenotyping of whole blood samples was carried out with three distinct flow cytometry panels to analyse myeloid, granulocyte and lymphoid subsets. (Figure 1A, Supplementary Table 3). Each panel was supplemented with counting beads to allow accurate assessment of cell counts. 19 of the 54 acute patients had paired plasma samples that permitted quantification of immune mediators by Luminex multiplex microbead-based immunoassay. The cohort was strongly biased towards males of which two patients had fatal outcomes (3.7%).

**Figure 1:**
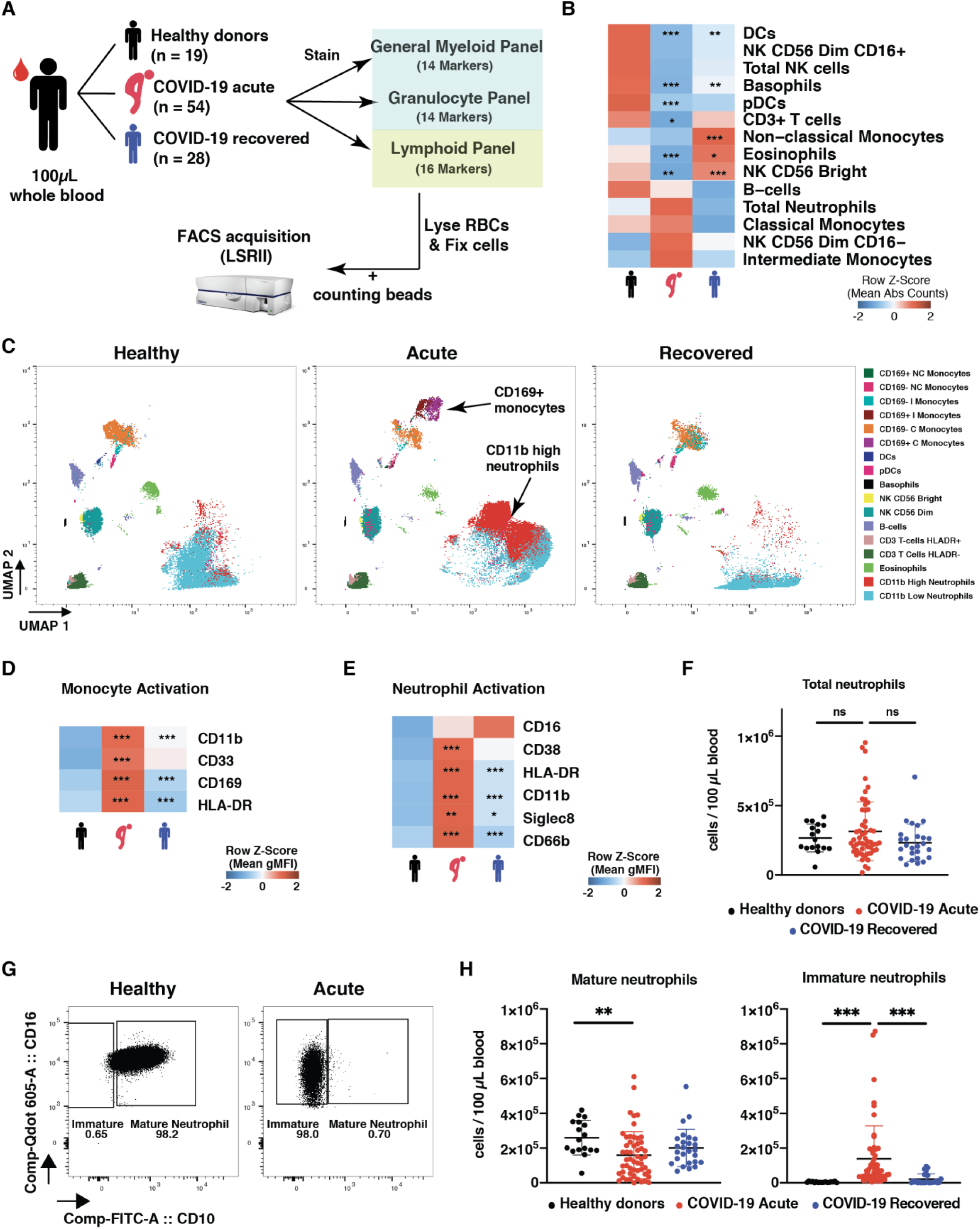
SARS-CoV-2 infection induces a decrease in immune cells in peripheral blood. (a) Schematic representation of flow cytometry workflow. (b) Heatmap representation of row z-score of mean absolute cell counts across the groups. Individual plots are shown in Supplementary Figure 1A. (c) UMAP clustering of CD45+ immune cells. (d) Heatmap representation of row z-score of monocyte activation markers mean geometric MFI (gMFI) across the groups. (e) Heatmap representation of row z-score of neutrophil activation markers mean geometric MFI (gMFI) across the groups. (f) Absolute neutrophil counts. (g) Representative plot of mature and immature neutrophil gating strategy in healthy control or acute COVID-19 patient. (h) Mature (CD10+) and Immature (CD10-) Neutrophil Abs counts. Absolute counts were analysed by Kruskal-Wallis using Dunn correction for multiple comparison, gMFI was analysed by Brown-Forsythe and Welch ANOVA using Dunnett T3 correction for multiple comparison. For heatmaps, stars shown in acute column represent healthy vs acute comparison. Stars shown in recovered column represent acute vs recovered comparison. ns non-significant. *p<0.05, **p<0.01, ***p<0.001

The FACS analysis revealed a declined cell count for eosinophils, basophils, total T-cells, dendritic cells (DCs), natural killer (NK) CD56 Bright, and plasmacitoid DCs (pDCs) in patients with acute COVID-19 infection (Figure 1B, Supplementary Figure 1A). No significant changes were observed for B-cells, total monocytes, and total NK cells (Figure 1B, Supplementary Figure 1A). Unbiased analysis by Uniform Manifold Approximation and Projection (UMAP) and graph-based clustering however identified with CD169^+^ monocytes and CD11b^high^ neutrophils, two additional clusters with high variation in acute patients (Figure 1C). Further analysis showed that the monocytes presented with an increased expression of CD169 (strong type I interferon signature marker ^14^), increased expression of CD11b and HLA-DR, as well as CD33, a constitutive PI3K signaling inhibitor ^15,16^ (Figure 1D, Supplementary Figure 1B).

Similar to the monocytes, neutrophils showed a significant upregulation of CD11b, CD66b, Siglec 8, CD38 and HLA-DR, suggesting that they were activated in response to SARS-CoV-2 infection (Figure 1E, Supplementary Figure 1C). Interestingly, despite this activation phenotype, an increase in the overall number of circulating neutrophils during acute SARS-CoV-2 infection based on conventional phenotypic markers (CD66b and CD16) was observed only in a small subset of our cohort (Figure 1F). However, in-depth analysis of neutrophil subsets allows discrimination between immature (CD16^low/high^CD10^−^) and mature (CD10^+^) subsets (Figure 1G)^17-19^. Overall, a significant increase of immature neutrophil numbers was observed in acute patients as compared to healthy donors or recovered patients, while the number of mature neutrophils decreased (Figure 1H).

### CD8 and γδ T-cell populations are the most affected lymphocyte subsets

To better characterise COVID-19-induced lymphopenia, levels of CD8, CD4, **γδ** (i.e. VD1 and VD2), and mucosal-associated invariant T-cells (MAIT, CD3^+^VA7.2^+^CD161^+^) were assessed during acute infection. Results showed a decrease in circulating MAIT, CD8^+^ and VD2 T-cells (Figure 2A). However, circulating VD1 T-cells did not vary in numbers, and CD4^+^ T-cells did not show a significant decrease during acute infection (Figure 2A). Interestingly, levels of regulatory T-cells (Treg) and CD4^+^CD161^+^ T-cells increased in recovered patients as compared to acute patients (Figure 2A).

**Figure 2:**
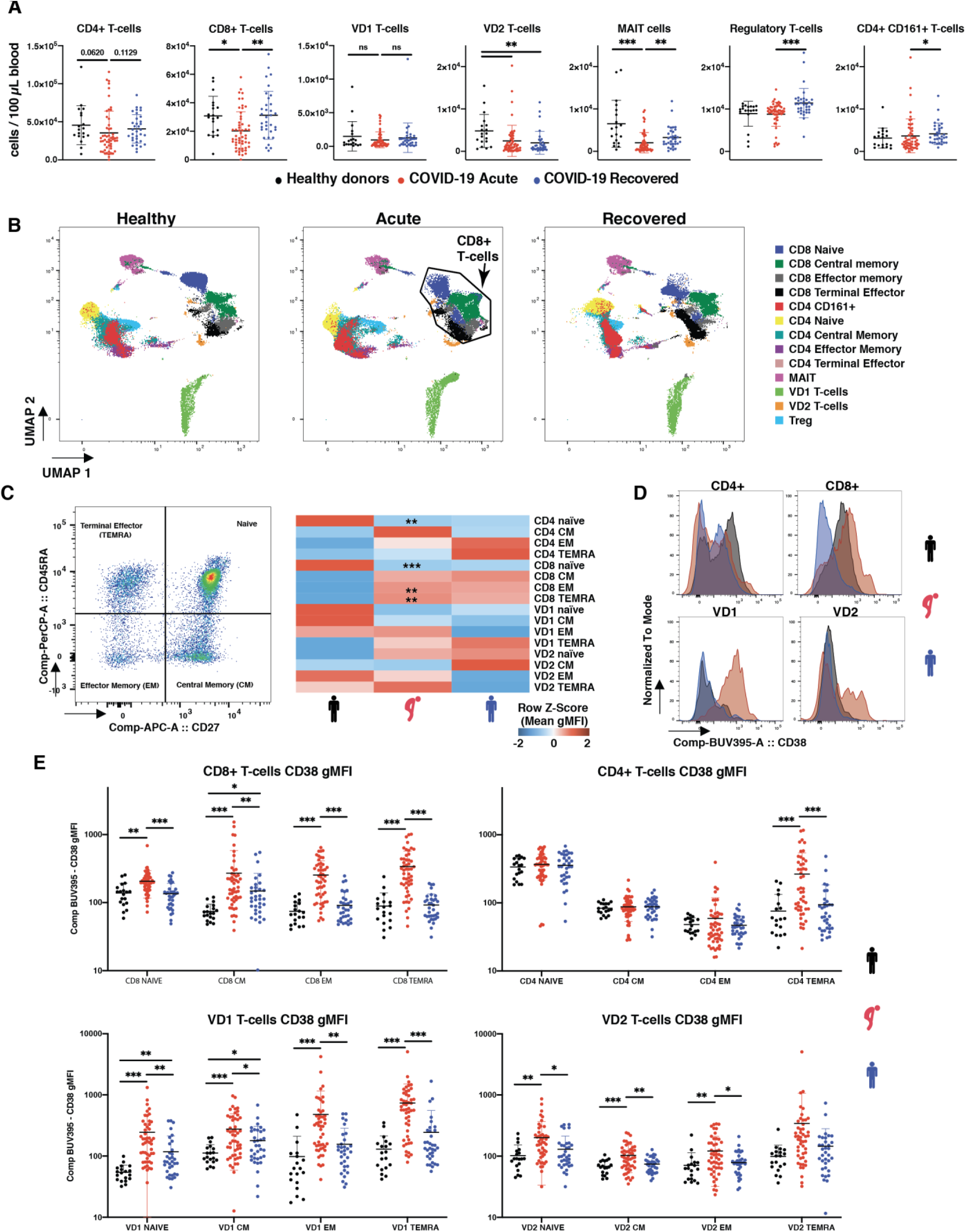
SARS-CoV-2 infection induces general lymphopenia and CD8, VD1 and VD2 activation. (a) Absolute counts of T-cell compartments in healthy donors, acute and recovered COVID-19 patient. (b) UMAP clustering of CD3+ cells. (c) left panel: CD45RA and CD27 gating strategy; right panel: heatmap representation of mean frequencies of T-cell differentiation across the groups, individual plots given in Supplementary Figure 2. (d) Representative histogram of CD38 expression in CD4, CD8, VD1 and VD2 T-cells. (e) Changes in CD38 gMFI in naïve, CM, EM and TEMRA for CD8, CD4, VD1 and VD2 T-cells. Absolute counts were analysed by Kruskal-Wallis using Dunn correction for multiple comparison, gMFI was analysed by Brown-Forsythe and Welch ANOVA using Dunnett T3 correction for multiple comparison. For heatmaps, stars shown in acute column represent healthy vs acute comparison. Stars shown in recovered column represent acute vs recovered comparison. *p<0.05, **p<0.01, ***p<0.001

Next, UMAP analysis was done on CD3^+^ cells to visualise changes in differentiation states within the T-cell compartments (Figure 2B). UMAP visualisation suggests that phenotypic modulation in the CD8^+^ cluster was the most pronounced during SARS-CoV-2 infection (Figure 2B). In order to validate this observation, CD45RA and CD27 markers were used to analyse the frequency of naïve (CD45RA^+^CD27^+^), central memory (CM, CD45RA^-^CD27^+^), effector memory (EM, CD45RA^-^CD27^-^) and terminal effector (TEMRA, CD45RA^+^CD27^-^) amongst the T-cell populations (Figure 2C, Supplementary Figure 2A). In agreement with the UMAP analysis, CD8^+^ T-cells showed a change in differentiation profile from naïve in favour of EM and TEMRA (Figure 2C). Noticeably, the frequency of naïve CD4^+^ T-cells decreased but was not reflected in a significant increase of a specific differentiated population (Figure 2C).

In addition, UMAP analysis also suggested changes in VD1 and VD2 populations that were not reflected in terms of differentiation (Figure 2B-C). Therefore, we investigated the expression of general activation marker CD38 (Figure 2D). In this context, we observed that all differentiation stages of CD8^+^ T-cells, VD1 and VD2, had higher expression of CD38 except VD2 TEMRA (Figure 2E). On the other hand, CD4^+^ T-cells only showed activation of the TEMRA compartment (Figure 2E). Together, our data suggest that while circulating cell counts were generally decreased for T-cells, SARS-CoV-2 differentially impacts the different T-cell subsets in terms cell counts, differentiation and expression of CD38.

### Granularity of clinical severity is reflected by immune cell counts

In order to associate the data with the clinical severity we separated the patients into four different groups: no pneumonia, pneumonia only, pneumonia and hypoxia, and pneumonia and hypoxia requiring ICU admission (Figure 3A) ^20,21^. This allowed estimation of cell counts in those groups and identification of markers that potentially depict disease severity. Consistent with previous studies on CD4 and CD8 lymphopenia ^6,22,23^, CD8^+^, CD4^+^, MAIT, VD1 and VD2 T-cells showed a gradual reduction in the peripheral blood with increasing disease severity (Figure 3B). The effect was more pronounced for CD8^+^ and VD2 T-cells (Figure 3B), suggesting a strong activation and infiltration of these cells in the lungs.

**Figure 3:**
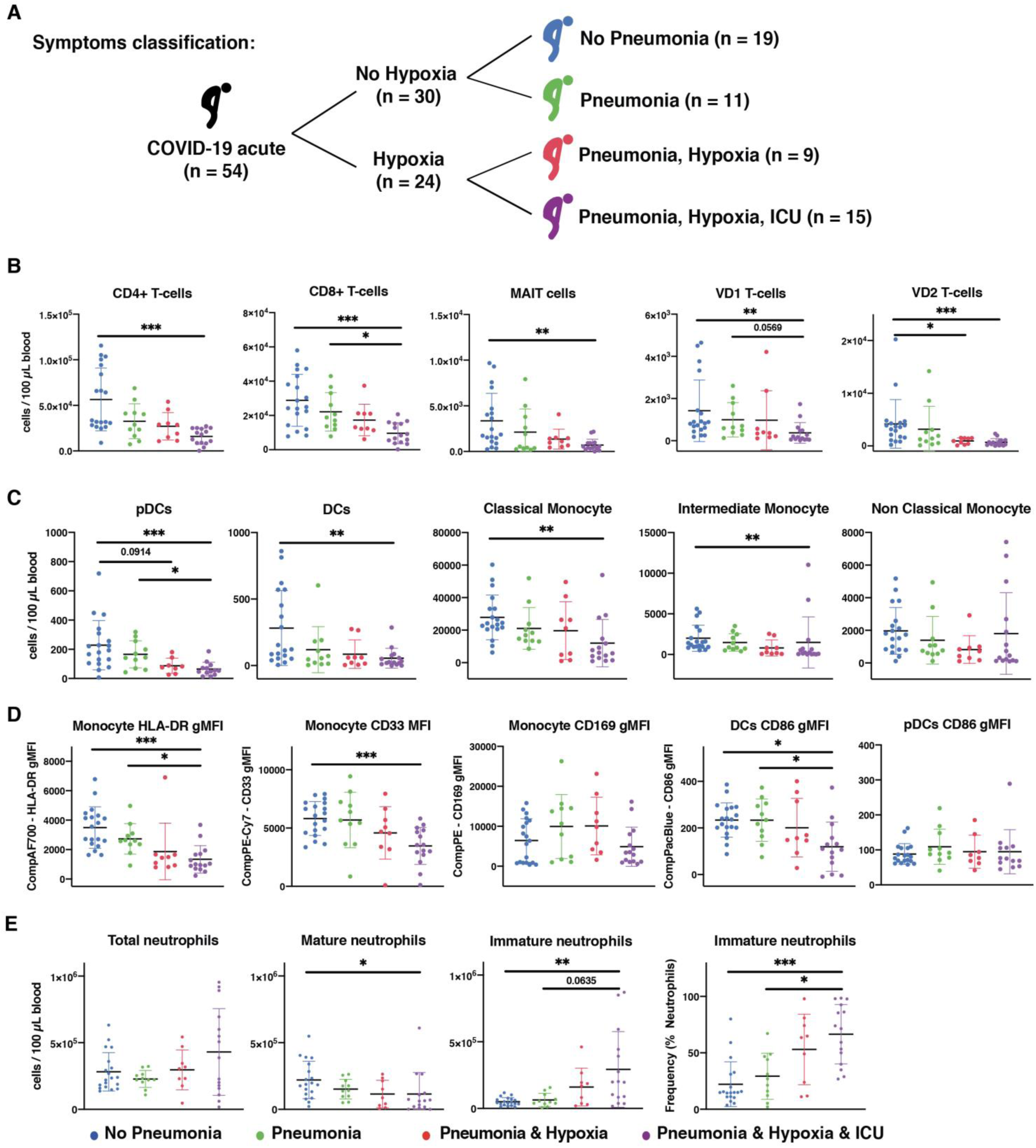
Patient symptoms are reflected in immune cell variations. (a) Schematic representation of clinical symptoms in the patient cohort. (b) Absolute counts of T-cells across the severity (c) Absolute counts of antigen presenting cells across the severity. (d) gMFI of activation markers on antigen presenting cells. (e) Absolute counts and frequency in neutrophil compartments. Absolute counts were analysed by Kruskal-Wallis with Dunn multiple testing correction, gMFI was analysed by Brown-Forsythe and Welch ANOVA with Dunnett T3 multiple testing correction. *p<0.05, **p<0.01, ***p<0.001

Cell counts in various myeloid subsets showed a similar decreasing profile with severity for pDCs, DCs, classical and intermediate monocytes (Figure 3C). In contrast to cell counts, myeloid activation markers showed differential trends with severity (Figure 3D). CD86 expression on DCs, HLA-DR and CD33 expression on monocytes followed a gradual decrease with increasing severity (Figure 3D). Expression of CD169 on monocytes was decreased in ICU patients, while CD86 expression on pDCs was consistent across severity groups (Figure 3D). Together, these results suggest that the remaining circulating monocytes and DCs in severe cases have a dysregulated phenotype.

While total circulating neutrophils showed no significant change with disease severity, neutrophilia was only observed in some patients with severe clinical complications (Figure 3E). Particularly, there was a change in the composition of neutrophil subsets in accordance to disease severity, where an increase in the immature neutrophil cell count and frequency was accompanied with a decrease of mature neutrophils (Figure 3E). These results suggest that immature neutrophils could reflect disease severity much more accurately than total neutrophil counts.

### Immature neutrophil absolute count correlates with cytokines

Neutrophil-to-Lymphocyte Ratio (NLR) or Neutrophil-to-CD8 T-cell Ratio (N8R) were proposed to be good diagnostic and prognostic markers for severe COVID-19 respiratory disease ^23,24^. However, these studies observed increased neutrophils in severe cases which was not consistent with our observations and in another study ^25^ (Figure 1F and 3E). To validate that the identified populations would be good markers of disease severity, a correlation analysis with analyte levels in available paired plasma samples was performed (Figure 4A, Supplementary Figure 3). Interestingly, strong correlation scores were observed between analytes and immature neutrophil counts (Figure 4A, Supplementary Figure 3A), rather than with total neutrophil counts (Figure 4A, Supplementary Figure 3B). The strongest correlations were observed between immature neutrophil counts and IL-6 (rho=0.6747, p=0.0015), and IP-10 (rho=0.7596, p=0.0002) (Figure 4B).

**Figure 4:**
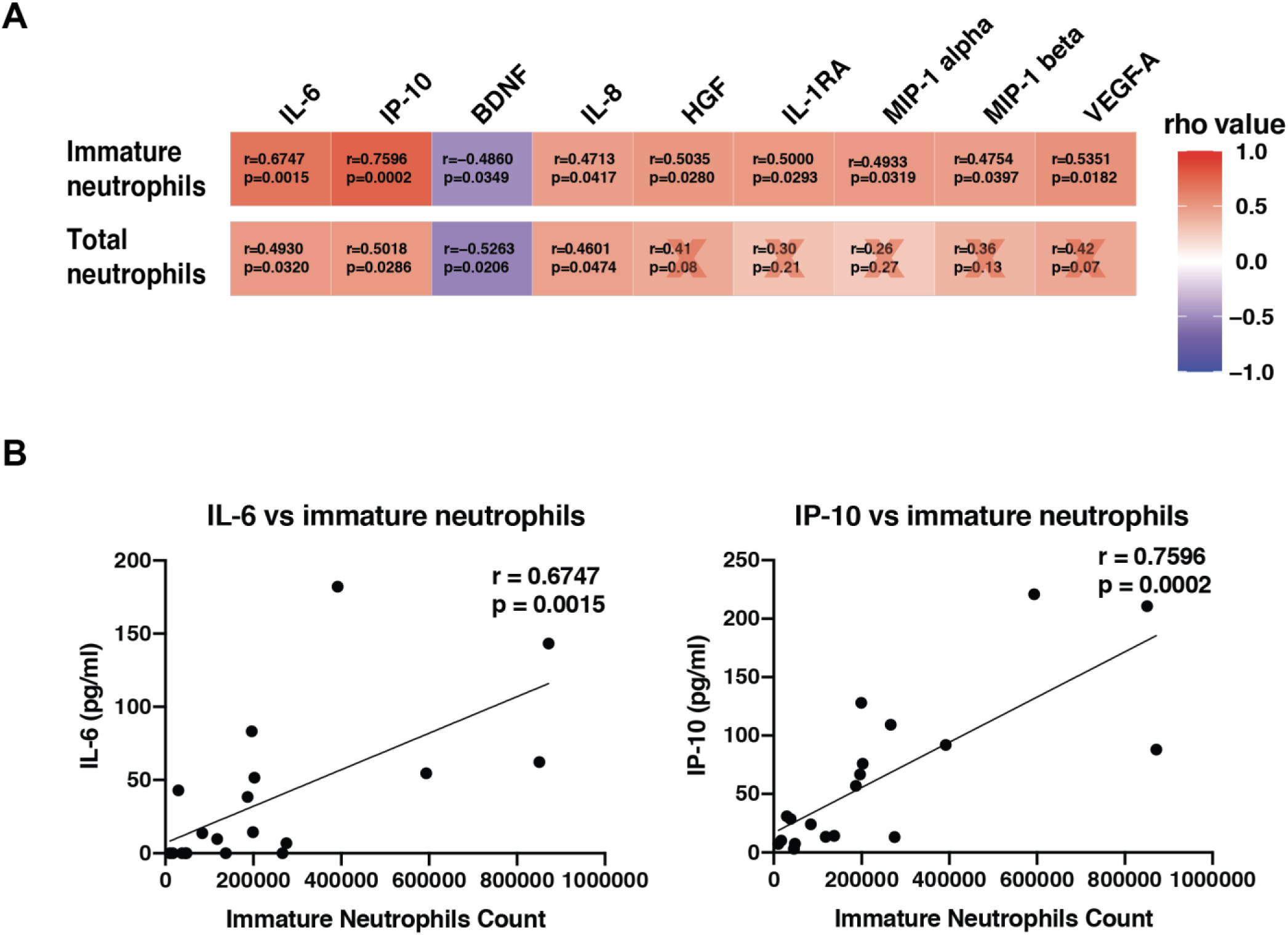
Immature neutrophils correlate with several analytes in paired patient plasma. (a) Spearman correlations between total neutrophils or immature neutrophils and plasma analytes. Red cross represents non-significant correlations. (b) Individual plots of Spearman correlations between immature neutrophil counts and IL-6 and IP-10. Line was drawn using simple linear regression.

In addition, strong correlations were also observed between mature neutrophils, monocytes and intermediate monocytes, as well as CD8 and VD2 T-cell counts (Supplementary Figure 3C). These results suggest that immature neutrophils counts can potentially be used as sensitive and reliable indicators of disease severity.

### Immature neutrophil to VD2 T-cell ratio as an improved prognostic marker

We next assessed if an immature neutrophil-to-CD8 T-cells ratio (iN8R) or VD2 T-cell counts ratio (iNVD2R) could be a better prognostic marker of disease severity as compared to the current proposed NLR and N8R ^23,24^. To differentiate patients with and without pneumonia, iNVD2R performed better than N8R or iN8R with an area under receiver operating characteristic (AUROC) curve of 0.8451 (95% confidence interval CI: 0.7379-0.9523) vs 0.806 (95% CI: 0.6911-0.9210) and 0.7158 (95% CI: 0.5754-0.8562) respectively (Figure 5A). In addition, to differentiate patients with and without hypoxia, an AUROC of 0.9111 (95% CI: 0.8306-0.9916) was obtained for iNVD2R as compared to 0.8931 (95% CI: 0.8044-0.9817) for iN8R and 0.7958 (95% CI: 0.6781-0.9136) for N8R. These results indicate that iNVD2R and iN8R could be good markers for severe respiratory disease.

**Figure 5:**
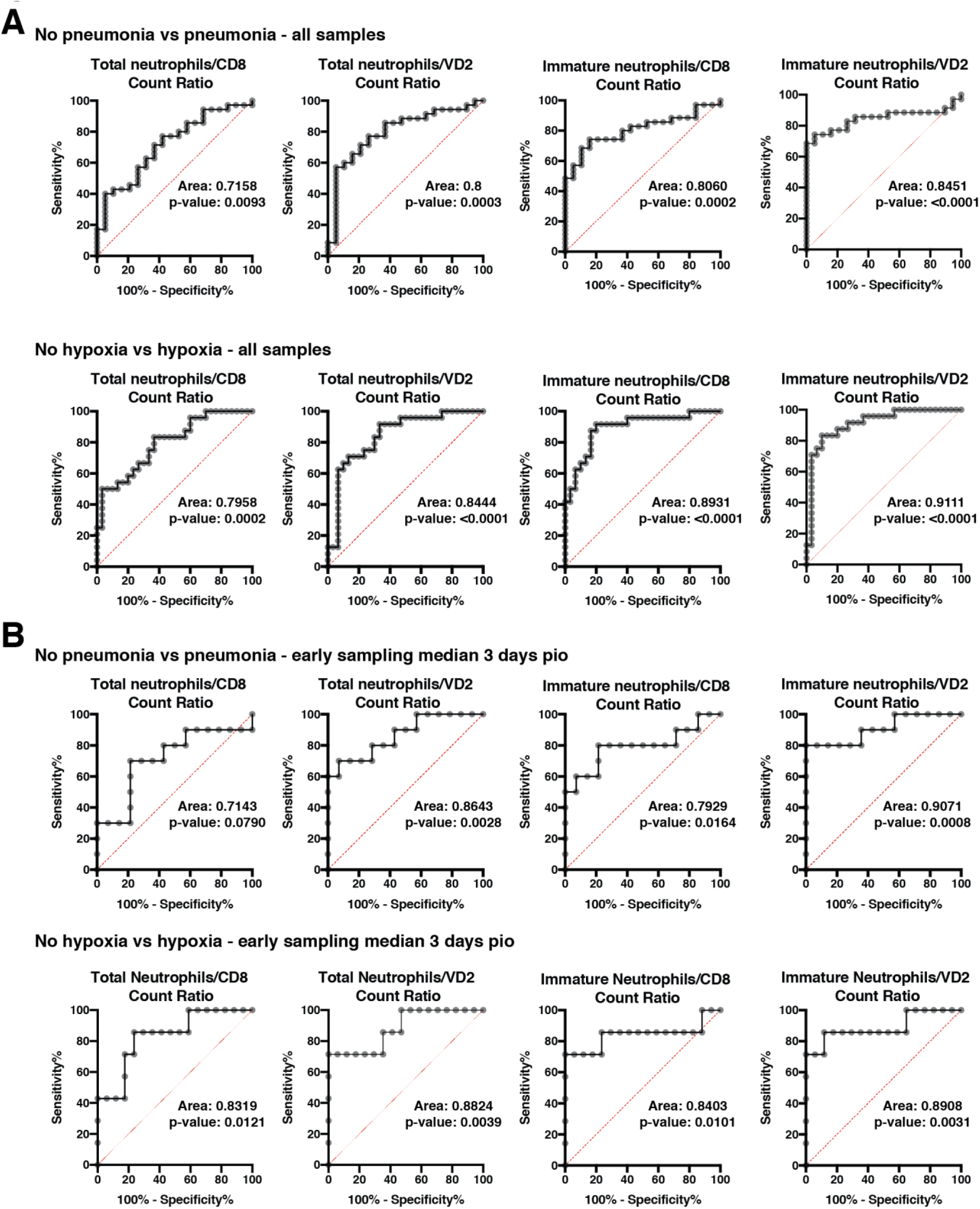
Immature neutrophil to VD2 T-cell ratio is an early prognosis marker for pneumonia and hypoxia symptoms. (a) ROC curve analysis comparison was performed for pneumonia and hypoxia symptoms between absolute counts of total neutrophils to CD8 T-cell ratio, total neutrophils to VD2 T-cell, immature neutrophils to CD8 T-cell ratio, and immature neutrophils to VD2 T-cell ratio. (b) Similar analysis was performed on a subset of 24 early samples taken up to 7 days pio with a median of 3 days pio. ROC curve was analysed using Wilson/Brown method. 95% confidence interval and standard error for panel B are given in Table 1.

To assess if this analysis could have predictive prognostic value in hospitalisation settings to improve patient management, we repeated the analysis with the samples that were acquired before 7 days pio (24 patients, median pio = 3 days). AUROC for iNVD2R showed strong prognostic value for pneumonia onset (0.9071) as well as for onset of hypoxia (0.8908) (Figure 5B, Table 1). Our data show that immature neutrophil counts are better in predicting disease severity as compared to total neutrophil counts. Importantly, they can be used in a ratio with CD8 or VD2 lymphocyte counts to improve the current N8R predictive ratio.

**Table 1:** ROC curve analysis for neutrophils to T-cell ratios in patients with pneumonia or hypoxia compared to those without as presented in Figure 5b.

| <u>Variable</u> | <u>Pneumonia</u> |  |  | <u>Hypoxia</u> |  |  |
| --- | --- | --- | --- | --- | --- | --- |
|  | AUC<br>(95% CI) | Std.Error | p-value | AUC<br>(95% CI) | Std.Error | p-value |
| Total neutrophils /<br>CD8 T-cells | 0.7143<br>(0.4909-0.9377) | 0.1140 | 0.0790 | 0.8319<br>(.6526-1) | 0.09149 | 0.0121 |
| Total neutrophils /<br>VD2 T-cells | 0.8643<br>(0.7135-1) | 0.07694 | 0.0028 | 0.8824<br>(.07239-1) | 0.08083 | 0.0039 |
| Immature neutrophils /<br>CD8 T-cells | 0.7929<br>(0.5884-0.9973) | 0.1043 | 0.0164 | 0.8403<br>(0.6079-1) | 0.1186 | 0.0101 |
| Immature neutrophils /<br>VD2 T-cells | 0.9071<br>(0.7754-1) | 0.06723 | 0.0008 | 0.8908<br>(0.7160-1) | 0.08915 | 0.0031 |
| ROC analysis was performed on COVID-19 patients between 2 to 7 days pio (24 patients, median 3 days pio). ROC curve was built by plotting true positive rate (sensitivity) against false positive rate (100%- sensitivity) and AUC was calculated from the plot. ROC, receiver operating characteristic ; AUC, area under curve ; CI, confidence interval ; Std.Error, standard error. |  |  |  |  |  |  |

## Discussion

In this study, immunophenotyping of peripheral blood from COVID-19 patients revealed a significant shift in the ratio between mature and immature neutrophils associating with severity. The increased numbers of immature neutrophils and the disappearance of mature neutrophils likely reflect gradual and sustained mobilisation of these cells into the lungs in response to an ongoing inflammation, leading to premature release of immature neutrophils from the bone marrow ^19^. Supporting this hypothesis, a recent study investigated several myeloid populations between circulating PBMCs and the lung lavage of COVID-19 patients showed that granulocytes represent up to 80% of total CD45^+^ lung infiltrates ^26^. In addition, autopsies of COVID-19 fatalities showed typical lesions associated with toxic neutrophil effects ^27,28^. In line with this observation, marked morphological abnormalities of the circulating neutrophils were reported in COVID-19 patients ^25^. These cells present typical hallmarks of immature neutrophils and their precursors such as band shaped nuclei and a lower expression of CD10 and CD16 ^29^. Consistent with our data, a recent study on a small number of patients reported that the presence of “low density inflammatory neutrophils” was strongly associated with disease severity and IL-6 levels ^30^. This CD11b^int^CD44^low^CD16^int^ low density neutrophil population is likely constituted primarily of CD10^−^ immature neutrophils.

In addition, immature neutrophil numbers strongly correlated with IL-6 and IP-10. IL-6 and IP-10 are consistently upregulated during a cytokine storm and are associated with severe ARDS ^9,10,31,32^. While some studies report inflammatory monocytes as the source of IL-6 ^9,33,34^, our results suggest that immature neutrophils could also be a non-negligible source of IL-6 during COVID-19-induced cytokine storm. Indeed, neutrophils have been found to produce biologically relevant amounts of IL-6 after engagement of TLR8, a toll like receptor recognising single strand RNAs of viral or bacterial origin ^35,36^. Since IL-17 operates upstream of IL-1 and IL-6, and is a major orchestrator of sustained neutrophils mobilisation ^37^, it is plausible that IL-17 could significantly affect the neutrophils compartment in COVID-19 patients. Consistent with this hypothesis, CD4 T-cells in COVID-19 patients are skewed towards a Th17 phenotype ^13^, and we also observed increased CD4^+^CD161^+^ T-cells in recovered patients. These CD4^+^CD161^+^ T-cells are known to be either IL-17 producer cells or their precursors ^38^. Thus, our results could reflect the re-circulation of these cells from the lung or secondary lymphoid organs after infection and support the possibility of IL-17 in mediating neutrophil damage to the lungs. Together, this would support proposed anti-IL-17 or JAK2 inhibitor therapies for severe COVID-19 disease ^39-41^.

In addition to the changes in the heterogeneity of neutrophils, a strong decrease in T-cells was observed, especially in subsets that possess cytolytic activity such as CD8, VD1 and VD2 T-cells. These results are consistent with other studies showing a decrease of CD8^+^ during COVID-19 disease ^12,13^. As for VD2 T-cells, which are not MHC-restricted T-cells ^42,43^, we showed a general decrease in the periphery with disease severity. This is in line with other inflammatory disease such as psoriasis ^44^ and Crohn’s disease ^45^. However, in the lungs, during chronic obstructive pulmonary disease, γδ T-cell counts have been reported to be significantly lower in induced sputum (IS) and bronchoalveolar lavage (BAL) but not in peripheral blood, suggesting unclear inflammatory mechanisms that could influence γδ T-cells counts in the periphery ^46^. Interestingly, γδ T-cells, in particular VD2, are known to participate in influenza immune response ^47^, and actively recruit and activate neutrophils to the site of infection or inflammation ^48,49^. Activated, neutrophils have also been found to inhibit γδ T-cells functional capacity, promoting the resolution of inflammation ^50,51^. Therefore, it will be essential to investigate the neutrophil to γδ T-cells relashionship present in lungs of SARS-CoV-2 infected patients.

During aging, VD2 T-cell counts in the periphery have been shown to decrease with age. Elderly individuals generally have systemic chronic low-grade inflammation, which we previously termed “inflamm-aging”, with higher basal levels of molecules such as CRP, TNF-a and IL-6 ^52,53^. These similarities in modulation of VD2 T-cell counts and cytokines between COVID-19 severity and aging could explain why elderly individuals are more susceptible to severe disease, since they have a higher basal level of inflammation and lower level of VD2 T-cells as compared to the young.

Our results indicate that an early post illness onset iNVD2R, accessible through a simple 5 colours flow cytometry panel (CD3; VD2; CD66b/CD15; CD10; CD45), would be an excellent prognostic screening tool for predicting probable patient progression to pneumonia or hypoxia. Moreover, CD8 could also be included in the flow cytometry panel as a fallback option since VD2 counts could be decreased by medication, such as Azathioprine, as well as underlying conditions, such as inflammatory bowel disease, aging or psoriasis, which could be risk factors for COVID-19 ^45^. Analysis of the proposed parameter would allow for a more accurate and earlier prognosis due to the interconnection between neutrophils and Vδ2 T cells, which can then be utilised for early therapeutic interventions, improve patient triage and better healthcare resource management.

## Material and Methods

### Study design

This was an observational cohort study of patients with PCR-confirmed COVID-19 who were admitted to the National Centre for Infectious Diseases, Singapore. All patients with COVID-19 in Singapore, regardless of the severity of infection, are admitted to isolation facilities until clinical recovery and viral clearance. Supportive therapy including supplemental oxygen and symptomatic treatment were administered as required. Patients with moderate to severe hypoxia (defined as requiring fraction of inspired oxygen [FiO_2_] ≥40%) were transferred to the intensive care for further management including invasive mechanical ventilation where necessary.

Sample Size: No power analysis was done. Sample size was based on sample availability. Randomization: No randomization was done. Blinding: Clinical parameters were made available after data analysis.

### Ethics statement

Written informed consent was obtained from participants in accordance with the tenets of the Declaration of Helsinki. For COVID-19 blood/plasma collection, “A Multi-centred Prospective Study to Detect Novel Pathogens and Characterize Emerging Infections (The PROTECT study group)”, a domain specific review board (DSRB) evaluated the study design and protocol, which was approved under study number 2012/00917. Healthy volunteers samples were obtained under the following IRB “Study of blood cell subsets and their products in models of infection, inflammation and immune regulation” under the CIRB number 2017/2806 from SingHealth (Singapore).

### Donor information

Patients who tested PCR-positive for SARS-CoV-2 in a respiratory sample from February to April 2020 were recruited into the study ^54^. Demographic data, disease onset date, clinical score and SARS-CoV-2 RT-PCR results during the hospitalisation period were retrieved from patient clinical records. Relevant information are given in Supplementary Table 1. Patients were classified in different clinical severity groups depending on the presence of pneumonia, hypoxia and the need for ICU hospitalisation. For healthy volunteers, demographic data are provided in Supplementary Table 2. Blood was collected in VACUETTE EDTA tubes (Greiner Bio, #455036) or Cell Preparation Tubes (CPT) (BD, #362753) and 100 μL of whole blood was extracted for each FACS staining panel (Supplementary Table 3).

### Multiplex microbead-based immunoassay

When available, plasma fraction was harvested after 20 minutes centrifugation at 1700 x *g* of blood collected in BD Vacutainer CPT tubes (BD, #362753). Plasma samples were treated by solvent/detergent treatment with a final concentration of 1% Triton X-100 (Thermo Fisher Scientific, #28314) for virus inactivation at RT for 2 hours in the dark under stringent Biosafety laboratory 2+ conditions (approved by Singapore Ministry of Health) ^55^. Immune mediator levels in COVID-19 patient plasma samples across acute samples were measured with by Luminex using the Cytokine/Chemokine/Growth Factor 45-plex Human ProcartaPlex™ Panel 1 (ThermoFisher Scientific, #EPX450-12171-901). Data acquisition was performed on FLEXMAP® 3D (Luminex) using xPONENT® 4.0 (Luminex) software. Data analysis was done on Bio-Plex ManagerTM 6.1.1 (Bio-Rad). Standard curves were generated with a 5-PL (5-parameter logistic) algorithm, reporting values for both mean florescence intensity (MFI) and concentration data. Internal control samples were included in each Luminex assay run to allow for detection and normalisation of plate-to-plate and batch-to-batch variation. A correction factor was obtained from the differences observed across the multiple assays with these controls and this correction factor was then used to normalise all the samples. Analyte concentrations were logarithmically transformed to ensure normality. Analytes that were not detectable in patient samples were assigned the value of logarithmic transformation of Limit of Quantification (LOQ).

### Flow cytometry

Whole blood was stained with antibodies as stated in Supplementary Table 3 (100 μL of whole blood per flow cytometry panel) for 20 minutes in the dark at RT. Samples were then supplemented with 0.5 mL of 1.2X BD FACS lysing solution (BD 349202). Final FACS lysing solution concentration taking into account volume in tube before addition is 1X. Samples were vortexed and incubated for 10 min at RT. 500 μL of PBS (Gibco, #10010-031) was added to wash the samples and centrifugated at 300 x *g* for 5 min. Washing step of samples were repeated with 1 mL of PBS. Samples were then transferred to polystyrene FACS tubes containing 10 μL (10800 beads) of CountBright Absolute Counting Beads (Invitrogen, #36950). Samples were then acquired using BD LSRII 5 laser configuration using automatic compensations and running BD FACS Diva Software version 8.0.1 (build 2014 07 03 11 47), Firmware version 1.14 (BDLSR II), CST version 3.0.1, PLA version 2.0. Analysis of flow cytometric data was performed with FlowJo version 10.6.1. Gating strategies for panels A, B and C are presented in Supplementary Figures 4, 5 and 6 respectively.

### Statistical analysis

Statistical analysis was performed using Prism 8 (Graph Pad Software, Inc). For comparisons of absolute cell counts or frequency, Kruskal-Wallis Test corrected with Dunn’s method was performed. For comparisons of geometric Mean Fluorescence Intensity (gMFI) between three or more independent groups, Brown-Forsythe and Welch ANOVA using Dunnett T3 correction for multiple comparison was performed. For correlation analysis, spearman rank correlation was performed. p-values < 0.05 for correlations, while adjusted p–values<0.05 for all the other comparisons were considered significant.

### Data analysis and UMAP visualisation

UMAP: Gated cells were manually exported using FlowJo (Tree Star Inc.). Samples were then used for UMAP analysis using cytofkit2 R Packages with RStudio v3.5.2 ^56^. Five healthy, six acute and four recovered patients were each concatenated to its respective groups and 100000 cells were analysed using the ceil method. Custom R scripts were used to generate Z-score and correlation heatmaps.

## Acknowledgements

Authors would like to acknowledge all the support received on this project from the Singapore Immunolgy Network (SIgN), LN lab members Chek Meng Poh and Anthony Torres Ruesta, and SIgN flow cytometry facility, especially Ivy Chay Huang Low. We would also like to thank the study participants who donated their blood samples to this project, and the healthcare workers caring for COVID-19 patients.

## Author contributions

GC, WX, IK conceptualised, designed the panels, acquired, analysed and interpreted the data, and wrote the manuscript. MYA processed the patient blood, stained and fixed the samples. YHC, SWF, KJP, BL, CYPL,SNA, NKWY, RSLC, WH, AA, acquired and analysed the data. EFB, SSWC, BEY, YSL and DCL designed and supervised sample collection. OR, LR, LGN, AL and LFPN conceptualised, designed, analysed and wrote the manuscript. All authors revised and approved the final version of the manuscript.

## Data availability

Data can be obtained upon reasonable request to the corresponding author.

